# Early salt shock uncouples shoot - root acclimation in *Lobularia maritima*

**DOI:** 10.64898/2026.04.20.719653

**Authors:** Carlos González-Cobo, Roser Tolrà, Gheith Charif, Eliana Bianucci, Mercè Llugany

**Author notes:** Author for correspondence: Roser Tolrà.

## Abstract

- Salt stress triggers overlapping osmotic and ionic effects. In halophytes, rapid acclimation can obscure how early responses are coordinated across organs. The use of facultative halophytes and salt shocks provides a useful framework to resolve this transition.
- We investigated the first 24 h of salt shock responses in the facultative halophyte *Lobularia maritima* by integrating physiological, ionomic, transcriptional and phytohormonal analyses to resolve organ- and time-dependent acclimation dynamics.
- Salt shock induced a rapid but transient osmotic effect, with shoot turgor recovery after 8 h. This recovery was associated with sustained osmotic adjustment, proline accumulation and increased Na^+^ levels in shoots. Conversely, photosynthetic impairment persisted beyond osmotic recovery. Salt exposure rapidly reshaped shoot and root ionomes and was associated with dynamic expression of *LmSOS1*, *LmNHX1*, and *LmHKT1*, consistent with coordinated Na^+^ partitioning. Oxidative responses diverged between organs; shoots maintained a stable oxidative state, while roots exhibited progressive loss of meristem viability. Abscisic acid (ABA) was strongly accumulated at all time points and emerged as the dominant regulator of early responses.
- These results show that early salt acclimation in *L. maritima* is rapid but spatially and functionally uncoupled, combining fast shoot osmotic adjustment with persistent photosynthetic constraints and increased root vulnerability.

## 2. Introduction

Salt stress imposes a major constraint on plant growth because it disrupts growth, water relations, nutrient homeostasis, and metabolism (van Zelm *et al*., 2020). Two interconnected components are usually distinguished: an early osmotic phase caused by the reduction in external water potential, and a later ionic component associated with Na^+^ accumulation and the ensuing nutritional and metabolic imbalance (Munns & Tester, 2008). In practice, these components are difficult to separate, particularly in tolerant species in which rapid acclimation may blur the temporal transition between them (Hariadi *et al*., 2011). Resolving how plants coordinate these early responses remains essential for understanding salt resilience.

Halophytes are especially informative systems in this context because they maintain growth under salinity levels that are inhibitory or lethal for most glycophytes. Their tolerance largely relies on the coordinated regulation of osmotic adjustment, ion transport, redox homeostasis, and hormonal signalling (Tolrà *et al*., 2025). Precisely because halophytes acclimate rapidly, many early responses to salinity may be missed under gradual salt treatments (Shavrukov, 2013; Hsieh *et al*., 2025). Salt shock therefore offers a useful experimental framework to resolve the initial dynamics of stress perception and acclimation where osmotic and ionic effects are imposed simultaneously (Skorupa *et al*., 2019; Dong *et al*., 2025).

Studies in halophytes have shown that the first 24 h after salt exposure are particularly informative for understanding stress adjustment (Hsieh *et al*., 2025). Early changes in Na^+^ and K^+^ partitioning, antioxidant activity, and stress signalling have been reported in several salt-tolerant species, indicating that tolerance depends not only on the magnitude of the response but also on its temporal coordination (Ellouzi *et al*., 2011; Nikalje *et al*., 2018; Li & Zhang, 2025). However, organ-specific dynamics during this early period remain poorly resolved, especially in facultative halophytes, which provide a useful system to understand how robust salt acclimation is achieved without the extreme specialization found in obligate halophytes (Ellouzi *et al*., 2014; Ludwiczak *et al*., 2023).

*Lobularia maritima* is a facultative halophyte in the Brassicaceae that has attracted interest because of its ecological breadth, ornamental value, and tolerance to salinity (Ben Hsouna *et al*., 2022; Koller *et al*., 2024; Yang *et al*., 2024). This species can withstand high NaCl concentrations and displays a shoot Na^+^ includer behaviour, together with osmoprotectant accumulation and enhanced antioxidant response under salt exposure (Ben Hsouna *et al*., 2020). Previous research has provided valuable information on its tolerance to prolonged salinity, but the transition between osmotic and ionic phases of salt stress remains unclear. We asked whether rapid recovery of plant water status is accompanied by equally rapid recovery of photosynthetic performance, how Na^+^-transport-related genes are associated with early ion partitioning, whether oxidative responses are similarly coordinated across organs, and which were the hormonal modifications that integrated the different salt and osmotic responses. To do so, we investigated the first 24 h of salt shock in *L. maritima* seedlings exposed to 400 mM NaCl combining physiological, ionomic, molecular, and hormonal analysis to resolve the temporal coordination of early salt responses in shoots and roots. Our results support a model in which early salt acclimation in *L. maritima* is spatially uncoupled, with rapid shoot osmotic adjustment occurring alongside sustained root vulnerability.

## 3. Materials and methods

### 3.1. Plant material and growth conditions

Seeds from a commercial variety of *L. maritima* (Vilmorin, Paris, France) were used in this study. Prior to sowing, seeds were surface sterilized by soaking in a sterilization solution (sodium hypochlorite [NaClO] 30% [v/v], 0.1% [v/v] Tween-20) for 15 min in constant agitation, followed by six washings with autoclaved 18 MΩ Mili-Q water, and left for 2 days submerged in GA_3_ 0.1 mM at 4 °C to stratify. Seeds were germinated in pots containing a mixture of peat and perlite (3:1), and seven-day old seedlings were transferred to individual hydroponic containers (50 mL) filled with ½ Hoagland nutrient solution (pH 5.7). Plants were grown under a 16 h light/8 h dark photoperiod, at 25 °C (day/night), 150 µmol m^-2^ s^-1^ light intensity, and 40% relative humidity, and distributed randomly among treatments and timepoints. The hydroponic solution was refreshed every three days to maintain constant nutrient concentrations. After seven days, seedlings (n=5) were transferred to ½ Hoagland solution supplemented with either 0 mM NaCl (Control) or 400 mM NaCl (Salt) for 1 h, 4 h, 8 h, 12 h and 24 h (ZT1, ZT5, ZT9, ZT13, and ZT24, respectively). Prior to harvesting, root architecture was analysed with WinRHIZO v.2009c software (RRID:SCR_017120). Plants were then harvested, shoot and root length and fresh weight were measured, and tissues were frozen in liquid nitrogen and stored at -80 °C until further use.

### 3.2. Plant water status

Relative water content (RWC) was determined according to the approach proposed by Barrs and Weatherley (1962). Plants were weighed immediately after harvesting (FW) and subsequently immersed in distilled water for 4 h for turgid weight (TW) measurement. Afterwards, the plants were oven-dried at 80 °C for 48 h to determine their dry weight (DW). The RWC was calculated as follows:

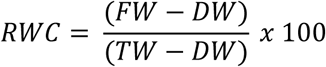

Leaf osmotic potential (Ψ_s_) was determined by measuring the freezing point depression (Furlan *et al*., 2012). Briefly, 0.1 g of frozen leaves were boiled in a water bath at 100 °C for 30 min in a 1.5 mL Eppendorf tube. The samples were then centrifuged at 20,800 × *g* for 10 min, and the supernatant was used to determine Ψ_s_ using an osmometer (Osmomat 3000, Gonotec®, Logan, USA). At least five biological replicates per treatment and time point were analysed.

### 3.3. Photosynthetic activity

Photosynthetic efficiency of *L. maritima* seedlings was determined *in vivo* from the second pair of leaves using a pulse-amplitude modulation (PAM) portable fluorometer (Mini-PAM II, Heinz Walz GmbH, Effeltrich, Germany). Measurements were performed in light-curve mode, following manufacturer’s instructions. Before measuring, plants were dark-adapted for 20 min. The following parameters were recorded: the indicator of PSII maximum yield (F_v_:F_m_), the effective quantum yield of PSII (Y[II]), the regulatory quantum yield of energy dissipation (Y[NPQ]), and the non-regulatory quantum yield of energy dissipation (Y[NO]). Five biological replicates were analysed per treatment and time point.

### 3.4. Shoot free proline content

Shoot proline content was measured following the method described by Bates *et al*. (1973). Briefly, 0.1 g of frozen leaves were homogenized in 2 mL of 3% sulfosalicylic acid. The solution was centrifuged at 10,600 × *g* for 10 min. The reaction mixture consisted of 1 mL of extract, together with 1 mL of ninhydrin mix (0.025% [w/v] ninhydrin, 60% [v/v] acetic acid, and 40% 6 M orthophosphoric acid [v/v]), and 1 mL of acetic acid. The mixture was incubated at 100 °C in a water bath for 1 h. The reaction was stopped by placing the samples in an ice bath for 20 min. The organic phase was separated by adding 2 mL of toluene, vortexing for 20 s, and collecting the upper organic phase. Absorbance was measured at 520 nm using toluene as a blank. Proline concentration was calculated from a standard curve generated with known concentrations of L-proline and expressed on a fresh weight basis.

### 3.5. Oxidative stress indexes and antioxidant enzymatic activity

Shoot hydrogen peroxide (H_2_O_2_) content was measured following the procedure described by Alexieva *et al*. (2001). Lipid peroxidation was analysed in *L. maritima* shoots by measuring the concentration of thiobarbituric-reactive substances (TBARS) (Heath & Packer, 1968).

The superoxide dismutase (SOD, EC 1.15.1.1) and catalase (CAT, EC 1.11.1.6) extracts were obtained by homogenizing shoot samples (0.1 g) with an extraction buffer (50 mM potassium phosphate buffer, pH 7.8, 0.5 mM EDTA, insoluble polyvinylpyrrolidone, and 0.5% [*v/v*] Triton X-100) using a mortar and pestle, followed by a centrifugation at 10,600× *g* for 10 min. Total protein content was measured according to Bradford assay, using a calibration curve containing of known concentrations of bovine serum albumin (Sigma).

SOD activity was determined according to Beauchamp & Fridovich (1971), following Bianucci *et al*. (2017) modifications. The reaction was performed by mixing 10 µL of protein extract to 290 µL of a reaction mix containing 777 µM of methionine, 75 µM of nitroblue tetrazolium (NBT), EDTA 0.54 µM, 8 µM riboflavin, and 50 mM phosphate buffer (pH 7.8). Samples were exposed to white light for 1 min, and absorbance was measured at 560 nm using a plate spectrophotometer (TECAN). The absorbance in darkness was used as blank. One unit of SOD activity was defined as the amount of enzyme required to inhibit the reduction of NBT by 50%. Three technical replicates were measured per sample.

CAT activity was quantified as described by Aebi (1984). Briefly, 980 µL of phosphate buffer (pH 7.4) was mixed with 10 µL 5 mM H_2_O_2_ and 10 µL of protein extract. The reaction was monitored at 240 nm in a UV–visible spectrophotometer (TECAN), measuring the decomposition of H_2_O_2_ for 180 s. One unit of CAT was defined as the amount of enzyme required to degrade 1 mmol of H_2_O_2_ per gram of FW. Three technical replicates per sample were measured.

### 3.6. Root viability assay

Membrane integrity of apical root meristems was evaluated using fluorescein diacetate and propidium iodide (FDA-PI) staining. Freshly harvested root tips were incubated for 3 min in FDA (12.5 μg mL^-1^), rinsed twice with Dulbecco’s Phosphate Buffered Saline (DPBS; 140 mM NaCl, 6 mM Na_2_HPO4, 4 mM KH_2_PO_4_, pH 7.4), and finally stained for 10 min in PI (5 μg mL^-1^). Excess PI was removed by washing with DPBS. Fluorescence was observed under an epifluorescence microscope using an emission filter of 520 nm and an excitation filter of 450-490 nm.

### 3.7. Mineral content

Plant material was oven-dried at 60 °C for 2 days and pooled until a final dry weight of 30 mg for shoots and 10 mg for roots. Samples and subsequently subjected to open-air predigested in Pyrex tubes using a 3:1 mixture of 69% HNO_3_ and 30% H_2_O_2_ overnight. Digestion was completed after 4 h at 110 °C using a hot-block digestion system (SC154-54-Well Hot Block, Environmental Express, Charleston, SC, United States). The concentrations of sodium (Na^+^), potassium (K^+^), calcium, (Ca^2+^), magnesium (Mg^2+^), sulphur (S), phosphorous (P), iron (Fe), manganese (Mn), zinc (Zn), and boron (B) were analysed in both shoots and roots using inductively coupled plasma optical emission spectroscopy (ICP-OES; Thermo Jarrell-Ash, Model 61E Polyscan, UK). For multivariate visualization, radial plots were used. Relative salt / control values were normalized as Z-scores calculated independently for each element.

### 3.8. Gene expression of Na^+^ transporters and related genes

Total RNA of about 100 mg of leaves and roots was extracted using Maxwell RSC Plant RNA Kit (Promega Biotech Ibérica SL, Alcobendas, Spain) following the manufacturer’s protocol. RNA concentration was quantified using a Nanodrop spectrophotometer (Thermo Fisher Scientific, Barcelona, Spain). One microgram of total RNA was used as a template to synthetize first-strand cDNA using the iScript cDNA Synthesis Kit (Bio-Rad, Barcelona, Spain) according to the manufacturer’s instructions. The cDNA was used as template for Reverse-Transcriptase quantitative real-time PCR (RT-qPCR) using iTaqTM Universal SYBR Green Supermix (Bio-Rad, California, United States). Detection of fluorescence emission was performed on a CFX384 Real-Time System (Bio-Rad, California, United States).

Sequences of the genes of the target genes *LmSOS1*, *LmSOS2*, *LmSOS3*, *LmNHX1*, *LmHKT1, LmUBQ10*-were identified from the *L. maritima* reference genome (Huang *et al*., 2020) using Blastn software based on the orthologous sequences of *A. thaliana*. Primers were designed using Primer-Blast and manually checked using Snapgene (San Diego, California, United States). The detailed description of the primers used is detailed in Table S1. Relative quantifications were performed using *LmUBQ10* as internal reference. Relative gene expression was calculated using the 2^-ΔΔCt^ method by Livak & Schmittgen (2001). Three biological replicates and three technical replicates per sample were included in each treatment and time-point analysed.

### 3.9. Leaf endogenous phytohormones

The concentrations of abscisic acid (ABA), jasmonic acid (JA), salicylic acid (SA), indole-3-acetic acid (IAA) and phenylacetic acid (PAA) were extracted and analysed according to Llugany *et al*. (2013). Briefly, 250 mg fresh *L. maritima* leaf powder was homogenized in an ice-cold mortar with 750 μL of extraction solution consisting of methanol:isopropanol:acetic acid (20:79:1, v/v) and a fixed concentration of deuterated internal standards. The supernatant was collected after centrifugation at 1000 × *g* for 5 min at 4 °C. These steps were repeated twice, and the combined supernatants were lyophilized. Samples were dissolved in 250 μL pure methanol and filtered with a Spin-X centrifuge tube filter of 0.22 μm cellulose acetate (Coastar, Corning Incorporated, Salt Lake City, USA). Phytohormones and their corresponding deuterated internal standards (ABA^2^-H6, IAA-d5, PAA-d5, JA-d5 and SA-d6) were purchased from Sigma-Aldrich (St. Louis, MO, USA). Chromatographic separation of plant samples and standards was carried out on a Luna Omega Polar C18 column, 2.1 x 150 mm ID and 1.7 μm particle size (Phenomenex, Torrance, CA, USA) using a UHPLC system (Agilent 1290 Infinity III, Agilent Technologies, Santa Clara, CA, USA) coupled to a triple quadrupole-linear ion trap mass spectrometer (QTRAP 7500, ABSciex, Framingham, MA, USA), operating in a negative electrospray ionization mode ESI (−). Quantification was performed using a calibration curve prepared with increasing amounts of external standards and a fixed concentration of a solution containing deuterated internal standards, by injecting extracted and spiked samples in multiple reaction monitoring (MRM) mode. Identification of phytohormones was based on the retention time and the presence of specific peaks in the MRM traces, as compared to those of the corresponding standards.

### 3.10. Data analysis

The data were analysed using RStudio (base R 4.3.3). Graphs were generated using the ggplot2 package (Wickham, 2016), complemented with ggprism visualization package (Dawson, 2025). Two-way ANOVA was used to test significant differences between means, considering *treatment* and *time point* independent variables (p-value < 0.05). For multiple comparisons, t-tests were applied for control vs. salt pairwise comparisons and Tukey HSD test for comparing group means across time points of the same treatment (p-value < 0.05).

## 4. Results

### 4.1. Osmotic readjustment is not accompanied by photosynthetic recovery under salt shock

Fourteen-day-old *Lobularia maritima* seedlings were exposed to a salt shock treatment consisting of 400 mM NaCl for 1, 4, 8, 12 and 24 h. Salt shock induced a rapid but transient osmotic effect in *L. maritima*, showing visible shoot wilting during the earliest stages of treatment (1-4 h; Fig. **1a**). This acute dehydration was supported by significantly lower RWC values in salt-treated plants at 1 h and 4 h compared with controls (Fig. **1b**). Despite the severity of the treatment, seedlings progressively regained shoot turgor, and leaf hydration was phenotypically restored by 8 h (Fig. **1a**). Consistent with this, RWC returned to control-like values from 8 h onwards, indicating that the osmotic impact of salt shock was largely confined to the first hours after exposure.

**Fig. 1.**
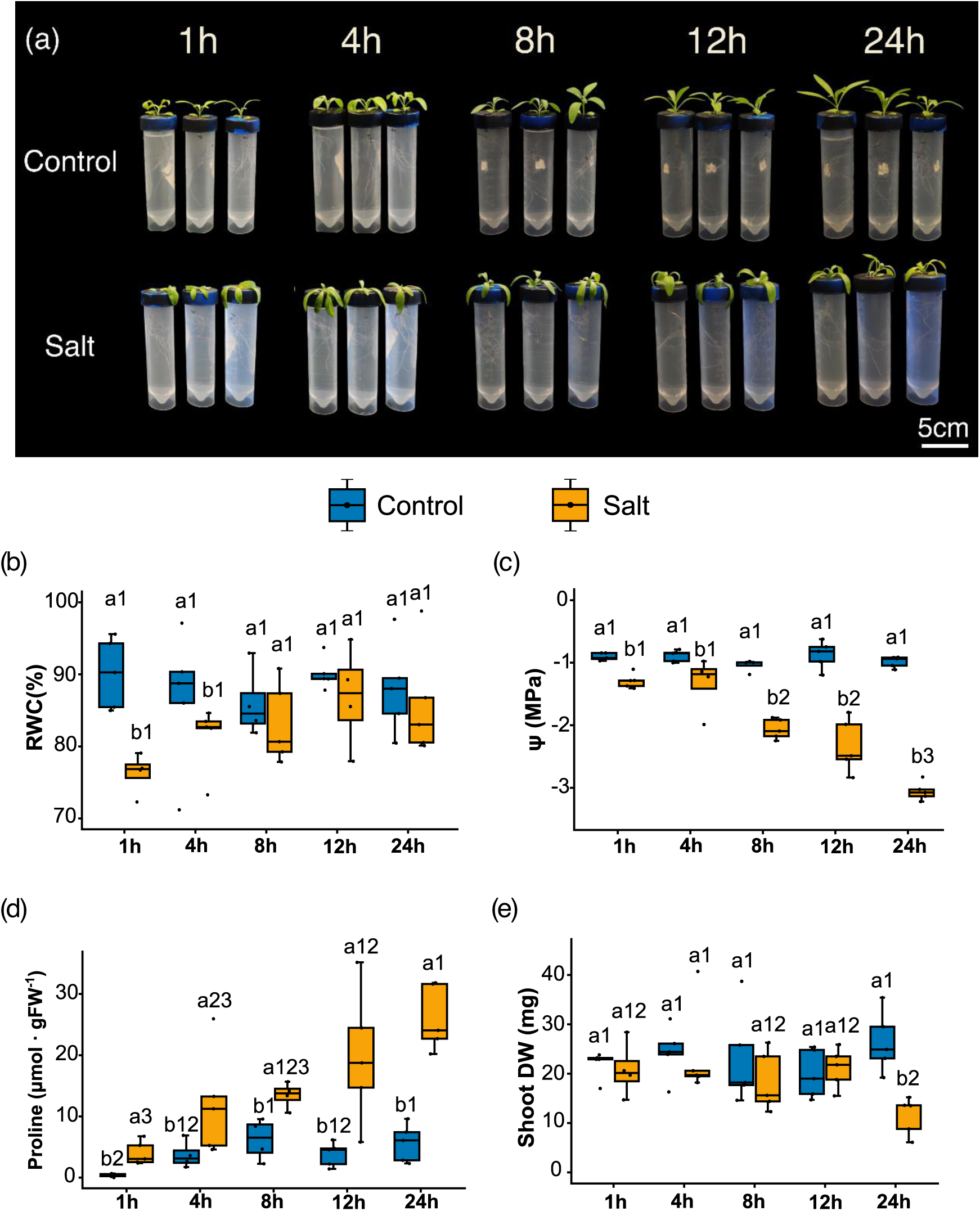
Phenotypic, growth, and osmotic status of *L. maritima* under salt shock. **(a)** Representative phenotypes of seedlings under control (C) or salt shock (S) conditions. **(b)** Shoot relative water content (RWC), **(c)** osmotic potential (Ψs), **(d)** free proline, and **(e)** dry weight in seedlings exposed to control or salt shock (400 mM NaCl) for 1, 4, 8, 12, or 24h. Different letters indicate significant differences among treatments within the same time point (t-test, *p* < 0.05); whereas different numbers indicate significant differences among time points at the same treatment (Tukey’s HSD, *p* <0.05).

This restoration of shoot water status was accompanied by a sustained decrease in leaf osmotic potential (Ψs), which was significantly reduced from the earliest time point analysed and reached -3.05 MPa after 24 h of salt exposure (Fig. **1c**). This osmotic adjustment correlates with a time-dependent accumulation of free proline in shoots, with salt-treated seedlings showing a progressive increase throughout the experiment and reaching a 4.6-fold increase compared to control plants at 24 h (Fig. **1d**). By contrast, growth-related penalties became apparent only at later stages. Shoot fresh weight was significantly reduced after 24 h of salt exposure with a 56.6% decrease relative to control seedlings (Fig. **1e**). Similarly, root growth alterations were only detected at the final time point, with reductions of 35% in root length and 40% in the number of forks (Table S1).

In contrast to water-status recovery, salt-treated seedlings showed a sustained impairment of PSII photochemistry and altered energy dissipation throughout the time course (Fig. **2**). Maximum PSII efficiency (Fv:Fm) decreased progressively under salt shock, with significant reductions relative to control plants at 1 h, 12 h and 24 h (Fig. **2a**). Similarly, Y(II) was significantly reduced from 4 h onwards (Fig. **2b**). Salt-treated plants also displayed higher non-photochemical quenching, Y(NPQ), during the first 12 h of treatment (Fig. **2c**). However, after 24 h of exposure, Y(NPQ) declined significantly, while non-regulated energy dissipation, Y(NO), increased significantly compared to control plants (Fig. **2d**).

**Fig. 2.**
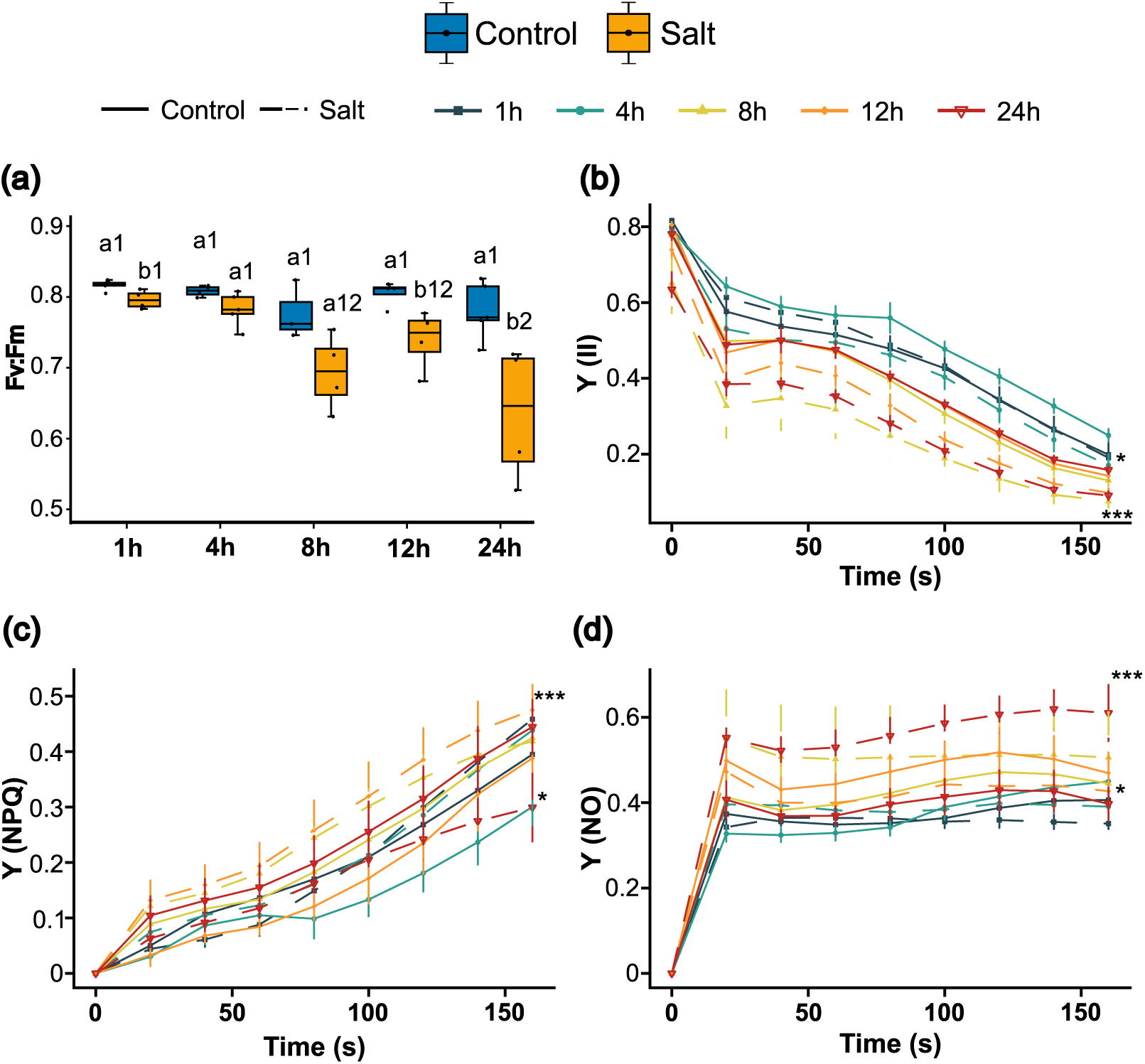
¡Error! No hay texto con el estilo especificado en el documento. **Photosynthetic alterations in *L. maritima* under salt shock**. **(a)** Maximum quantum efficiency of photosystem II (Fv:Fm) and time course measurements of **(b)** effective quantum yield of PSII (Y[II]), **(c)** non-photochemical regulatory quantum yield of energy dissipation (Y[NPQ]), and **(d)** non-regulatory quantum yield of energy dissipation (Y[NO]) in seedlings exposed to control or salt shock (400 mM NaCl) for 1, 4, 8, 12, or 24h. For Fv:Fm, different letters indicate significant differences among treatments at the same time point (t-test, *p* < 0.05), and different numbers indicate significant differences among time points within the same treatment (Tukey’s HSD, *p* <0.05). For Y(II), Y(NPQ), and Y(NO), asterisks indicate significant differences between treatments at the same time point (MANOVA, \**p* < 0.05; \*\*\**p* < 0.001).

### 4.2. Salt shock rapidly reshapes shoot and root ionomes

Salt shock caused a rapid and extensive reconfiguration of the ionome in both shoots and roots of *L. maritima* (Fig. **3a,b**). Significant differences between control and salt-treated plants were detected for nearly all elements analysed in at least one of the time points analysed, except for shoot Zn and root B, indicating that the early response to salinity extended beyond Na^+^ accumulation alone.

**Fig. 3.**
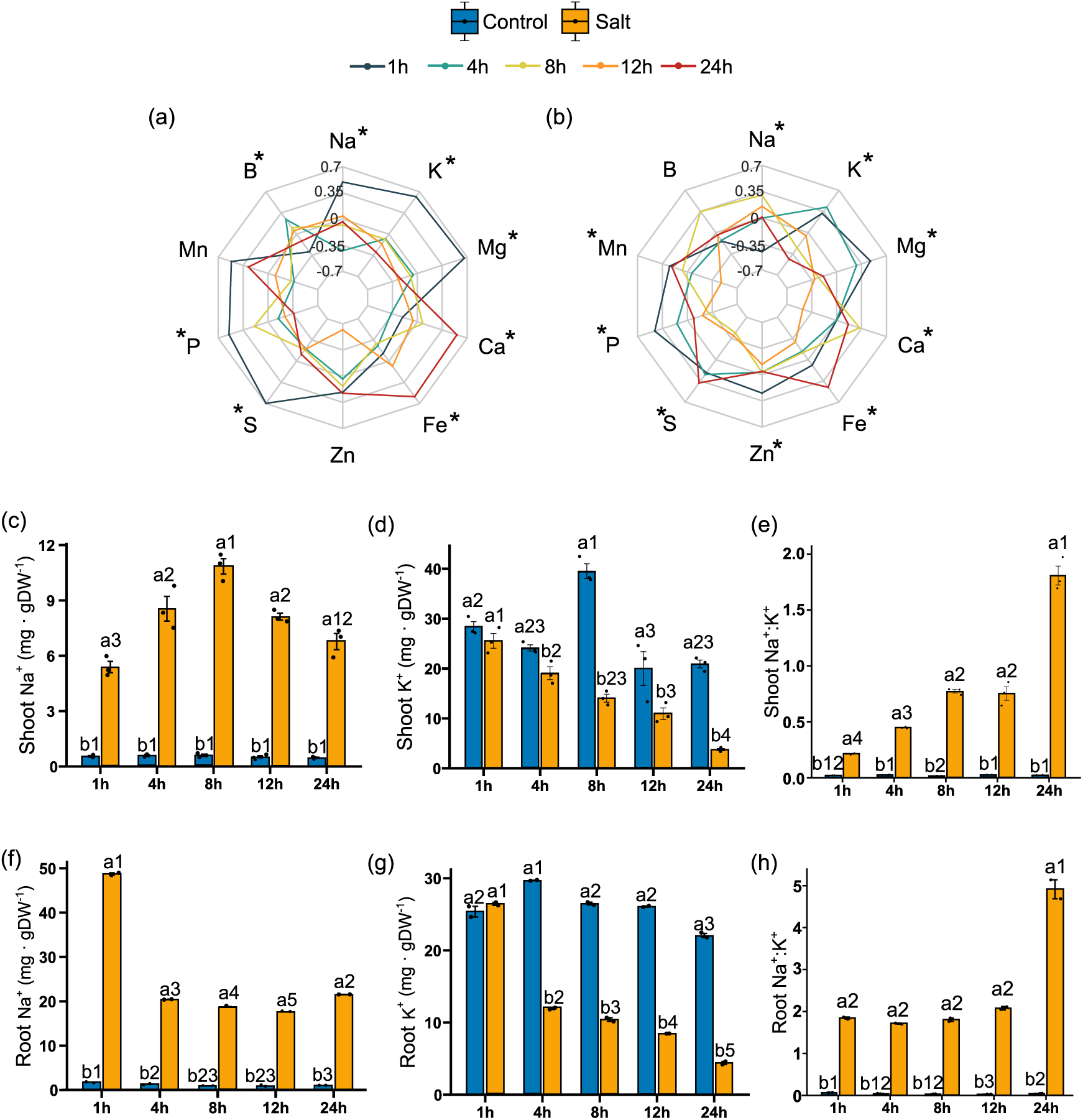
Changes in ionome homeostasis during salt shock. Radial plots showing relative *Lobularia maritima* ion concentrations in **(a)** shoots and **(b)** roots. **(c)** Shoot Na^+^, **(d)** K^+^, and **(e)** Na^+^:K^+^ ratio; **(f)** Root Na^+^, **(g)** K^+^, and **(h)** Na^+^:K^+^ ratio. Measurements were performed in seedlings exposed to control or salt shock (400 mM NaCl) for 1, 4, 8, 12, or 24 h. For radial plots, axis display Z-scores calculated independently for each element. Elements exhibiting significant differences at least at one time point (Tukey’s HSD, *p* < 0.05), based on the raw data, are indicated with an asterisk. For bar plots, different letters indicate significant differences among treatments at the same time point (t-test, *p* < 0.05), and different numbers indicate significant differences among time points within the same treatment (Tukey’s HSD, *p* <0.05).

As expected, Na^+^ accumulation was one of the earliest and most pronounced responses to salt shock. In shoots, Na^+^ content increased rapidly and reached a maximum at 8 h, followed by a slight decline at 12 h and 24 h (Fig. **3c**). In roots, by contrast, Na^+^ content peaked as early as 1 h (50 mg DW^-1^) and remained relatively stable thereafter (Fig. **3f**), indicating that Na+ partitioning between organs was dynamically regulated during the early response. In parallel, K^+^ suffered a time-dependent decrease in both shoots and roots over time (Fig. **3d,h**). Consequently, the Na^+^:K^+^ ratio increased progressively in shoots (Fig. **3e**), whereas in roots it remained relatively stable (2-fold increase compared to control) up to 12 h and increased sharply only after 24 h (Fig. **3i**). Notably, this late increase in roots was primarily associated with a marked decline in K^+^ rather than with additional Na^+^ accumulation.

The homeostasis of the other major cations, Ca^2+^ and Mg^2+^, was also altered under salt shock (Fig. **3a,b**). In shoots, Ca^2+^ content remained comparatively stable, except for a reduction at 12 h, while Mg^2+^ decreased significantly from the 8 h onwards (Fig. **S1a,b**). In roots, Ca^2+^ concentration was consistently lower under salinity across all time points, although some temporal fluctuations were observed (Fig. **S1a**). Root Mg²⁺ also declined over time and showed a more pronounced reduction than that observed in shoots (Fig. **S2b**). As a result, the Na^+^:Ca^2+^ ratio increased in both organs, although the shift towards Na^+^ dominance was more pronounced in roots than in shoots. By contrast, the Ca^2+^:Mg^2+^ ratio remained relatively stable in shoots except at the final time point, whereas it increased progressively in roots (Fig. **S3a,b; S4a,b**).

Macronutrient and micronutrient homeostasis were also affected by salt shock. In shoots, P and S levels were lower at 8 h and 12 h, and this trend was maintained in P at 24 h, although not significantly (Fig. **S1c,d**). In roots, reductions in these macronutrients were more evident and occurred earlier, with significant decreases already detected from 4 h onwards (Fig. **S2c,d**), suggesting a rapid impairment of nutrient uptake under salinity. Additionally, several micronutrients also displayed altered dynamics. In shoots, Fe accumulated markedly after 24 h of salt exposure, whereas in roots the concentrations of Fe, Zn and Mn were generally lower, although with some variation across ions and time points (Fig. **S1e S2e-g**). Accordingly, Mn:Fe and Zn:Fe ratios remained relatively stable in leaves until 24 h, when they declined under salinity (Fig. **S3c,e**). In roots, similar but earlier decreases were observed, with alterations in the Mn:Fe ratio detected from 4 h and in Zn:Fe from 12 h onwards. On the other hand, the Fe:S ratio, used as an indicator of Fe-S clusters necessary for photosynthesis and respiration (Astolfi *et al*., 2021), increased under salinity in both organs, with a stronger effect in roots than in shoots (Fig. **S4e,d**).

### 4.3. Oxidative responses diverge between shoots and roots under salt shock

Given the sustained impairment of photosynthetic performance and the rapid accumulation of Na^+^ in shoots, oxidative status was analysed in *L. maritima* seedlings exposed to salt shock. Salt exposure induced a dynamic but tightly regulated oxidative response in shoots (Fig. **4**). Shoot H_2_O_2_ levels increased significantly only after 8 h of salt treatment, whereas at 12 h and 24 h levels were lower than those detected in control plants (Fig. **4a**). Conversely, lipid peroxidation, estimated as TBARS content, showed a rapid and transient increase at the earliest stages of treatment, with significant peaks detected at 1 h and 4 h, followed by attenuation from 8 h onwards (Fig. **4b**). To assess whether this pattern was associated with antioxidant activation, the activities of key ROS-scavenging enzymes were quantified. SOD activity showed an oscillatory pattern over time, with higher values than controls at all time points except at 4 h (Fig. **4c**). CAT activity, by contrast, remained low in control plants but was strongly induced under salt shock, particularly during the early phase of treatment, peaking at 4 h with a 36-fold increase relative to control seedlings (Fig. **4d**). In contrast to the regulated oxidative response observed in shoots, root apical tissues showed progressive signs of cellular damage during salt shock (Fig. **4e-j**). FDA-PI staining of primary root meristems revealed cumulative membrane damage and loss of cell viability over time, with increasingly pronounced staining of non-viable cells at later time points.

**Fig. 4.**
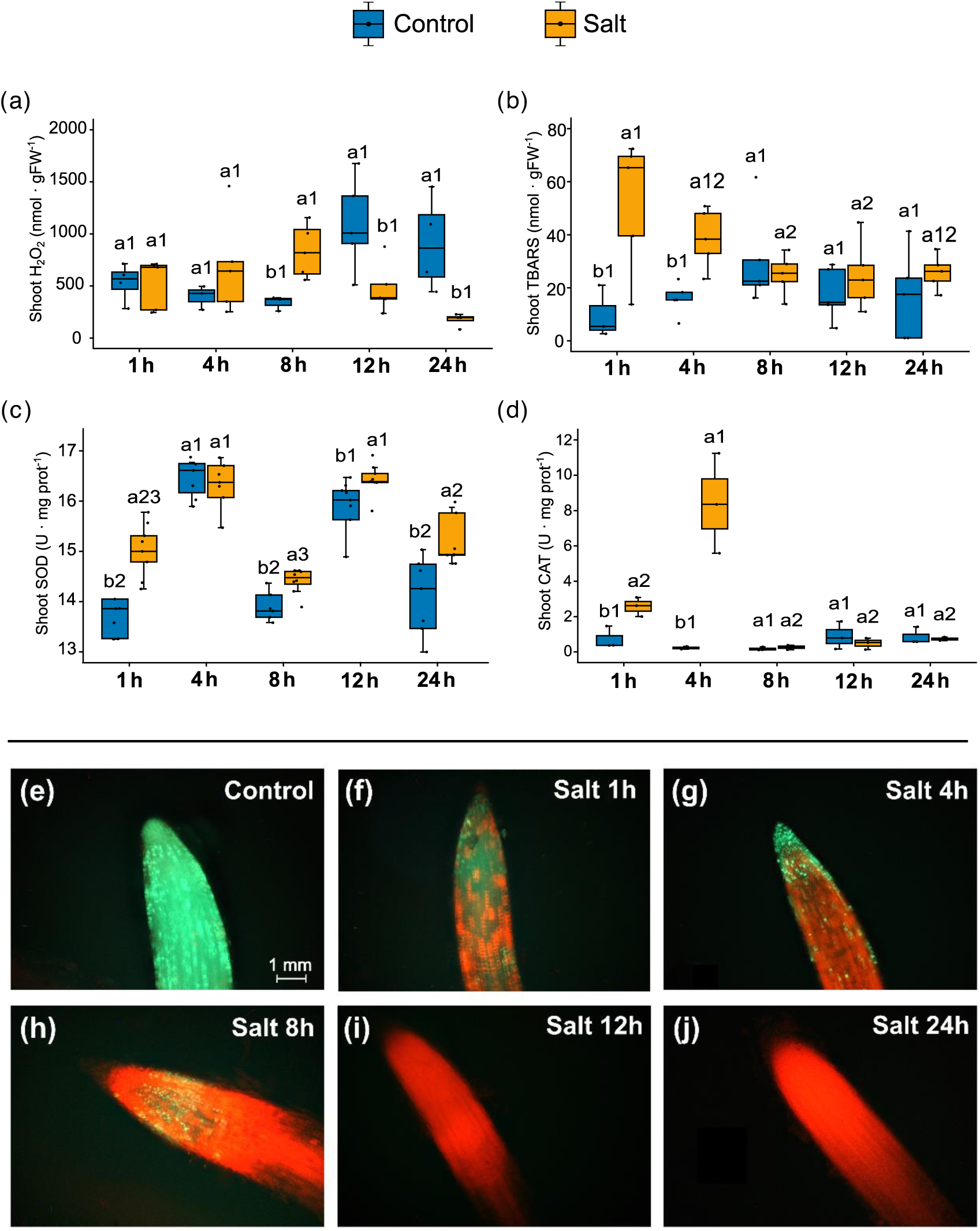
Oxidative stress status. Shoot **(a)** hydrogen peroxide, **(b)** thiobarbituric acid reactive substances, **(c)** SOD activity, and **(d)** CAT activity of *L. maritima* seedlings exposed to control or salt shock (400 mM NaCl) for 1, 4, 8, 12 or 24 h. (E-J) Root vital staining. FDA/PI staining of *L. maritima* primary root apical meristems exposed to control conditions **(e)** or salt shock (400 mM NaCl) for **(f)** 1 h, **(g)** 4 h, **(h)** 8 h, **(i)** 12 h, or **(j)** 24 h. For A-D, Different letters indicate significant differences among treatments at the same time point (t-test, *p* < 0.05), and different numbers indicate significant differences among time points within the same treatment (Tukey’s HSD, *p* <0.05). For E-J, one representative root per treatment and time point is shown.

### 4.4. Na^+^ transporter gene expression is dynamically coordinated during early salt shock

The contrasting temporal patterns of Na^+^ accumulation in shoots and roots suggested that early salt acclimation in *L. maritima* involved a dynamic regulation of Na^+^ transport processes (Fig. **3c,f**). To assess whether these ionomic changes were associated with transcriptional regulation of major Na^+^ transport-related genes, the expression of *LmSOS1*, *LmNHX1* and *LmHKT1* was analysed in shoots and roots during the first 24 h of salt shock. Due to the relevance of SOS pathway in salinity tolerance, the expression of the regulatory components *LmSOS2* and *LmSOS3* was also quantified.

Among the genes analysed, *LmSOS1* showed the strongest and most rapid transcriptional response to salt shock (Fig. **5a**). Transcript levels increased markedly in both tissues, with shoot expression reaching approximately 20-fold higher values than in control plants. This strong induction contrasted with the expression pattern of *LmSOS2*, which was generally reduced under salinity from 4 h onwards in both tissues, except in shoots at 24 h, where expression returned to control-like levels (Fig. **5b**). In turn, *LmSOS3* displayed a clear tissue-specific response: its expression was strongly induced in shoots at 4 h and 8 h, reaching 4.82- and 8.74-fold increases, respectively, whereas in roots it was significantly repressed after 4 h of salt exposure (Fig. **5c**). These contrasting dynamics show that while the Na^+^ transporter *SOS1* was enhanced, the transcription of their regulators differed markedly between shoots and roots during the early stages of salt shock.

**Fig. 5.**
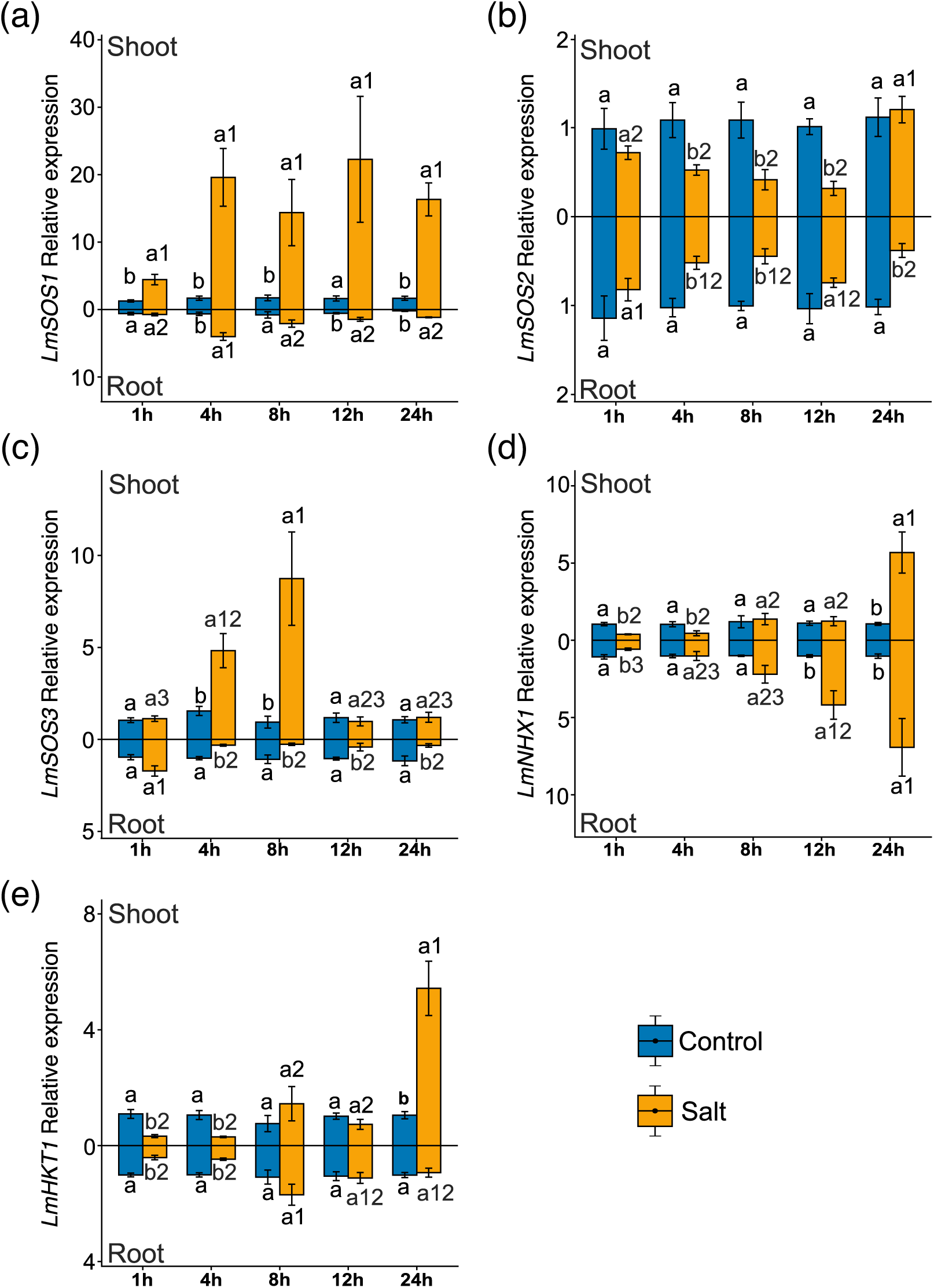
Expression of main Na^+^ transporters and regulators under salt shock. Relative expression levels of **(a)** *LmSOS1*, **(b)** *LmSOS2*, **(c)** *LmSOS3*, **(d)** *LmNHX1*, and **(e)** *LmHKT1* in shoots (upper panels) and roots (lower panels) of *L. maritima* seedlings exposed to control or salt shock (400 mM NaCl) for 1, 4, 8, 12 or 24 h. Different letters indicate significant differences among treatments at the same time point (t-test, *p* < 0.05), and different numbers indicate significant differences among time points within the same treatment (Tukey’s HSD, *p* <0.05).

The vacuolar Na^+^/H^+^ antiporter *LmNHX1* and the Na^+^ uniporter *LmHKT1* showed similar temporal patterns. In both cases, transcript abundance was reduced during the earliest stages of treatment, with *LmNHX1* downregulated at 1 h in both tissues and at 4 h in shoots, and *LmHKT1* repressed in both shoots and roots during the first 4 h of salt exposure. This early repression was followed by a later induction phase, particularly in shoots, where *LmNHX1* increased 5.68-fold and *LmHKT1* 5.42-fold after 24 h (Fig. **5d,e**).

### 4.5. ABA dominates the early shoot hormonal response to salt shock

To integrate the early physiological and molecular responses of *L. maritima* to salt shock, endogenous shoot concentrations of major phytohormones were quantified during the first 24 h of treatment (Fig. **6**). Among the hormones analysed, ABA levels showed the strongest and most consistent response to salinity. ABA levels were significantly higher than in control plants at all time points, with an induction of at least 25-fold and a maximum accumulation detected 4 h after salt exposure (Fig. **6a**). Notably, this peak coincided with the attenuation of the early osmotic effects observed in salt-treated seedlings (Fig. **1a,b**), supporting that ABA accumulation was closely associated with the osmotic acclimation phase.

**Fig. 6.**
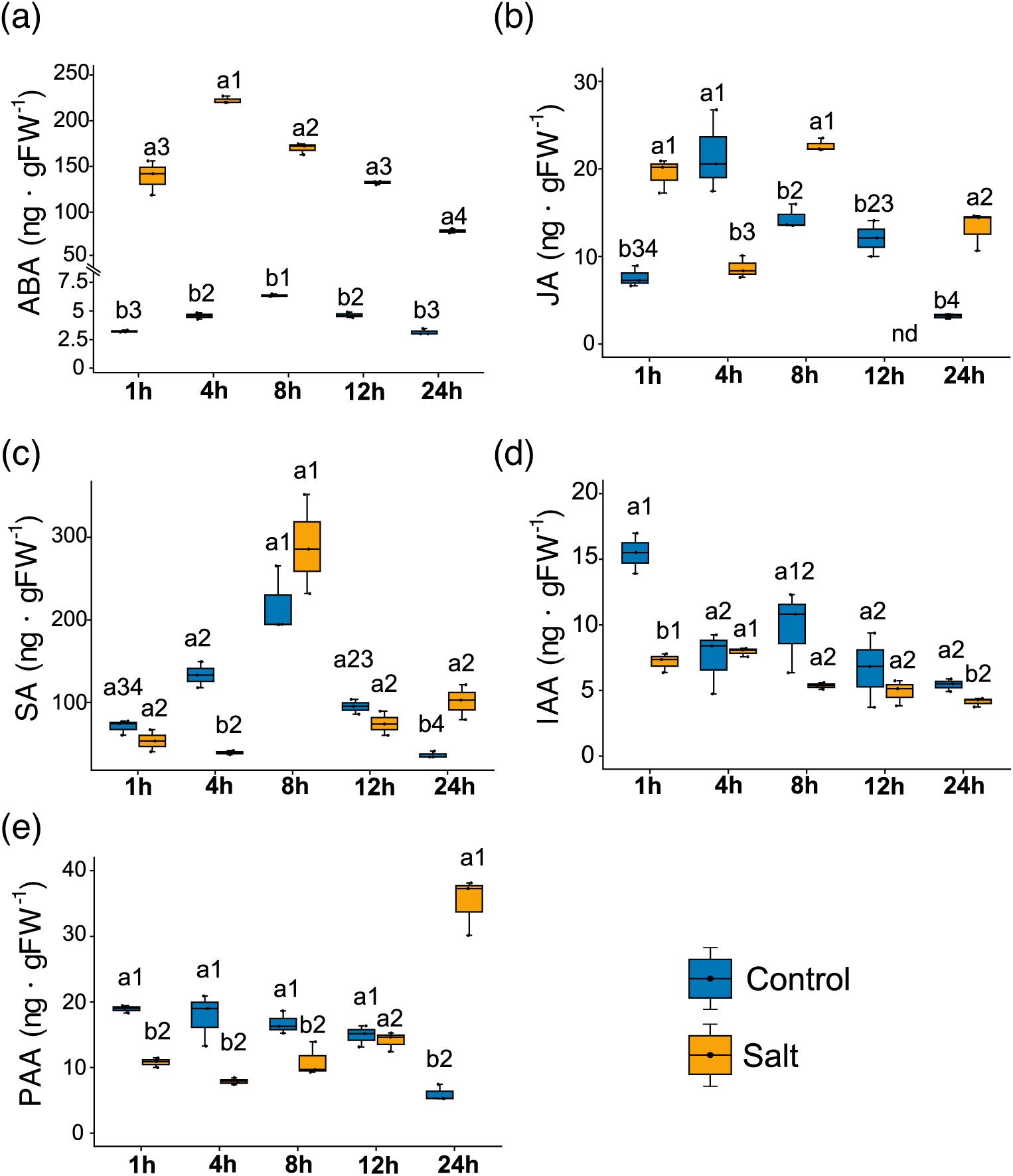
Shoot endogenous phytohormone levels under salt shock. **(a)** Abscisic acid, **(b)** jasmonic acid, **(c)** salicylic acid, **(d)** indole-3-acetic acid and, **(e)** phenylacetic acid in *L. maritima* seedlings exposed to control conditions or salt shock (400 mM NaCl) for 1, 4, 8, 12, or 24 h. Different letters indicate significant differences among treatments at the same time point (t-test, *p* < 0.05), and different numbers indicate significant differences among time points within the same treatment (Tukey’s HSD, *p* <0.05).

By contrast, JA displayed a more dynamic and oscillatory pattern. Under control conditions, JA levels peaked at 4 h, whereas under salt shock the temporal profile was altered, showing an opposite trend to that observed under control conditions (Fig. **6b**). SA content also fluctuated over time, with a maximum under control conditions at 8 h. Under salinity, however, SA levels decreased significantly at early time points, particularly at 4 h, and subsequently increased towards the end of the experiment, exceeding control values at 24 h (Fig. **6c**). Auxin-related responses were comparatively moderate. IAA showed a slight overall decreasing trend under salt shock, with significant reductions detected only at 1 h and 24 h (Fig. **6d**). In contrast, PAA levels decreased during the first 8 h of treatment and then progressively increased, ultimately reaching values above those of control plants at 24 h (Fig. **6e**).

## 5. Discussion

Although the temporal differentiation of osmotic and ionic effects of salt stress has been previously described (Munns, 2002; Shavrukov, 2013), the transition between these phases in salt-tolerant species remains scarcely studied. The great tolerance of halophytes to salinity makes them interesting models to understand how they modulate the responses at different organs and time points, and eventually to provide alternative salt tolerance mechanisms. In this study, we investigated the temporal and organ-specific responses against salt shock of the facultative halophyte *L. maritima*.

A central feature of the early response of *L. maritima* to acute salinity was its ability to rapidly restore shoot water status despite the severity of the treatment. Salt shock induced a transient dehydration phase, as evidenced by early shoot wilting and reduced RWC, yet both phenotypic turgor and leaf water status were largely restored within 8 h (Fig. **1a,c**). This pattern aligns with previous observations in halophytes**¡Error! Marcador no definido.**, in which hydration follows the initial osmotic disturbance imposed by external salinity (Ellouzi *et al*., 2011; Ryu & Cho, 2015; Méndez-Alonzo *et al*., 2016). Rather than avoiding the initial osmotic impact, *L. maritima* appears to cope with salt shock by rapidly compensating for it, indicating that osmotic restoration is an early and central component of its salt resilience.

This recovery was associated with a sustained decline in leaf osmotic potential together with progressive proline accumulation, supporting the establishment of active osmotic adjustment during the first hours of salt exposure. Although compatible solutes such as proline contribute to osmotic balance and act as osmoprotectants, inorganic ions are the primary determinants of cell turgor under salinity (Munns *et al*., 2020b). The observed reduction in shoot Ψ_s_ therefore likely reflects the combined effect of Na^+^ accumulation and osmolyte production, including proline (Fig. **1e****, S1a**). In addition to its osmotic role, proline may also contribute to stress tolerance through chaperone-like functions and redox buffering (Kant *et al*., 2006; Ji *et al*., 2022; Homayouni *et al*., 2024), suggesting a dual role in both water balance and cellular protection.

The temporal dynamics of ABA further supports this interpretation. ABA levels increased markedly and peaked at 4 h, coinciding with the recovery of the initial osmotic effects (Fig. **1a****, 6a**). Given the established role of ABA in regulating stomatal closure, osmotic adjustment and stress-responsive metabolism (Ryu & Cho, 2015), this timing suggests that ABA is closely linked to the reestablishment of water balance in *L. maritima.* The concomitant accumulation of ABA and proline is also consistent with ABA-dependent regulation of proline biosynthesis under abiotic stress (Savouré *et al*., 1997; Sharma & Verslues, 2010), and agrees with previous observations in *L. maritima* and other halophytes (Popova & Golldack, 2007; Ben Hsouna *et al*., 2020).

Notably, this early restoration of shoot water status did not prevent downstream growth penalties, which became evident only after prolonged exposure (Fig. **1e**). This temporal sequence indicates that hydraulic stabilization precedes the full recovery of physiological function. Although the precise transition between osmotic adjustment and complete recovery cannot be resolved with the present sampling resolution, the data clearly place this transition within the first hours following salt shock.

However, this rapid hydraulic recovery was not paralleled by a comparable restoration of photosynthetic performance. The persistent impairment of PSII photochemistry indicates that it remained functionally constrained beyond the period of acute dehydration (Fig. **2**), consisting with photoinhibition (Murata *et al*., 2007). These observations suggest that restoration of tissue hydration alone is insufficient to prevent downstream impairment of the photosynthetic apparatus under salt shock.

The decoupling between water balance and photosynthetic function likely reflects the transition from an initial osmotic phase to a later ionic and metabolomic phase of salt stress. Na^+^ accumulation in shoots can disrupt chloroplast function through effects on electron transport, thylakoid integrity and photochemical efficiency, thereby promoting photoinhibition (Allakhverdiev & Murata, 2008; Murata *et al*., 2007; Wang *et al*., 2024). The early increase in Y(NPQ) observed is consistent with a transient enhancement of regulated energy dissipation as an immediate protective response. However, the subsequent decline in Y(NPQ), together with the increase in Y(NO), indicates that this protective capacity was not fully sustained, leading to less controlled dissipation of excess energy.

The oxidative data agree with this interpretation. In shoots, H₂O₂ and lipid peroxidation showed transient or tightly regulated dynamics, while SOD and CAT activities were rapidly activated (Fig. **4a,c**). These changes in the oxidative status may represent a signalling threshold triggering the transition to ionic-phase responses (Ben Rejeb *et al*., 2015). In this regard, proline accumulation, in combination with SOD and CAT activities (Fig. **1d****, 4c-d**), may have contributed to this redox buffering (Renzetti *et al*., 2024). Additionally, the oscillatory SOD activity observed may have been caused by a circadian clock regulation of SOD-encoding genes (Jiménez *et al*., 2021). This suggests that, despite persistent limitations in PSII performance, *L. maritima* maintained a relatively controlled redox state in aerial tissues during the first 24 h of salt exposure. Similar patterns have been described in other halophytes, where photosynthesis performance is often affected under high salinity even when oxidative damage remains partially contained (Debez *et al*., 2008; Ellouzi *et al*., 2011; Yan *et al*., 2020). However, under certain conditions, salinity has also been reported to support the maintenance of photosynthetic function and redox balance, highlighting the context-dependent nature of these responses (Wali *et al*., 2016). Overall, these findings indicate that early acclimation is only partial: shoot water balance is restored rapidly, whereas photosynthetic function recovers more slowly, highlighting chloroplast performance as a particular sensitive component of the early salt response, especially under salt shock.

A second defining feature of the response of *L. maritima* to salt shock was its marked spatial asymmetry between shoots and roots. While shoots rapidly regained water balance and maintained a comparatively regulated oxidative state, roots exhibited stronger disruption of ion homeostasis together with progressive loss of meristem viability (Fig. **3,4**). These observations indicate that acclimation to acute salinity is not uniformly distributed across the plant but instead involves a pronounced functional uncoupling between organs.

The ionomic data reinforces this view. Although salt shock rapidly reshapes the shoot and root ionomes of *L. maritima*, affecting not only Na^+^ and K^+^ but also the broader mineral network, especially in roots (Fig. 3, Fig S1b-d, S2b-d). The reduction in K^+^ and Mg^2+^ may also contribute to the impaired photosynthesis observed under salt shock (Fig. 2). On the one hand, K^+^ role in photosynthesis involves the regulation of H^+^ efflux into the lumen, thereby maintaining ATPase activity (Bose *et al*., 2017). On the other hand, the reduction of foliar Mg^2+^ may reflect an impaired chlorophyll biosynthesis (Jamali Jaghdani *et al*., 2021) The reduction of root Ca^2+^, a key element in the activity of Ca^2+^ dependent kinases such as SOS3 (Ali et al., 2023; Su et al., 2020), could also contribute to explain the reduced expression of this gene in *L. maritima* salt-shock seedlings. The increased Na^+^:K^+^ and Na^+^:Ca^2+^ ratios in roots indicate a more severe ionic imbalance (Fig. 3h, S4a). In contrast, shoots appear to operate under a more buffered internal environment. Given that roots are the first organ exposed to external salinity, this pattern suggests that they act as an early site of ionic perturbation, potentially contributing to the modulation of shoot responses during the initial phase of salt exposure through the production of ROS and the controlled loading of ions (Gul *et al*., 2026).

At the cellular level, root meristems showed progressive membrane damage and loss of viability. K^+^ imbalance may have contributes to this deterioration, as K^+^ efflux has been linked to programmed cell death in salt-stressed roots (Wu *et al*., 2018). Primary roots are also known to be more sensitive than lateral roots under severe salinity (Ambastha *et al*., 2020). Notably, the effects observed here were more severe than those previously reported in *L. maritima* seedlings exposed to salt stress under acclimatory conditions (González-Cobo *et al*., 2024), suggesting that root meristem collapse under salt shock may reflect the limited capacity of these tissues to adjust when salinity is imposed abruptly. Meanwhile, the comparatively controlled oxidative dynamics observed in shoots suggest that aerial tissues are better able to contain oxidative perturbation despite impaired photosynthesis. Similar shoot-root divergence has been reported in other halophytes (Ellouzi *et al*., 2014). Altogether, these observations indicate that salt resilience in *L. maritima* involves a differential allocation of stress burden, with shoot function stabilized early while root tissue remain more vulnerable.

Early acclimation in *L. maritima* also involved dynamic regulation of Na^+^ transport and stress signalling. Although the species behaved as a Na^+^ includer, the temporal patterns of Na^+^ accumulation suggest an actively regulated process rather than passive ion loading. The transient peak of shoot Na^+^ and its stabilization in roots indicate an early phase of translocation followed by redistribution and intracellular management, consistent with the use of Na^+^ as an osmoticum during early acclimation (Munns *et al*., 2020a).

Gene expressions patterns support this interpretation. The strong and rapid induction of *LmSOS1* suggests a pivotal role in early Na^+^ transport, whereas the delayed activation of *LmNHX1* points to later vacuolar sequestration (Fig. **5a,d**). Similarly, early repression of *LmHKT1* in roots may favour shoot accumulation, while its later induction in shoots could contribute to the partial decline in leaf Na^+^ (Fig. **5e**). These dynamics indicate a coordinated balance between Na^+^ transport, retention and compartmentation, and aligns with previous observations in other halophytes (Katschnig et al., 2015). Interestingly, the canonical SOS regulatory module did not show fully coordinated expression. The inhibition of *LmSOS2* and *LmSOS3* contrast with previous findings and suggest that additional regulatory mechanisms may be involved (J. Huang et al., 2024; Liu et al., 2000; Rahman et al., 2021). Alternative pathways, including phospholipase D-and kinase-mediated signalling have been implicated in salinity responses (Quan *et al*., 2007; Ghars *et al*., 2012; Ji *et al*., 2013; Li *et al*., 2023).

Within this framework, ABA appears as a central regulator of early acclimation. By contrast, JA, SA and auxins displayed more variable dynamics, suggesting a more context-dependent contribution during the early response (Fig. **6b-e**). The circadian oscillations of JA and SA are consistent with observations in *A. thaliana*, with JA and SA peaks around mid-day and midnight, respectively (Goodspeed *et al*., 2012; Zhang *et al*., 2019). Although auxins can be produced ubiquitously, long-distance shoot-to-root auxin transport is required for proper regulation of root architecture, lateral root emergence, and halotropism (Blakeslee *et al*., 2019). The decreased activity of IAA is therefore consistent with the reduced number of root branches observed after 24 h of salt shock (Table S1).

## 6. Conclusions

This study provides a temporally resolved characterization of early salt shock responses in *L. maritima*. Exposure to 400 mM NaCl induced a rapid but transient osmotic stress, followed by restoration of water status through osmotic adjustment. Despite this recovery, salt shock severely altered photosynthetic performance, indicating that restoration of water balance does not imply full functional recovery. Salt exposure also triggered marked ionomic reconfiguration, with stronger disruption and progressive loss of viability in roots, revealing a pronounced spatial asymmetry in early responses. The temporal dynamics of Na^+^ accumulation, together with the expression profiles of key transporters, support coordinated regulation of Na^+^ partitioning, while divergence within the SOS regulatory components suggests additional layers of control for a fine-tune modulation of the response. At the hormonal level, ABA emerged as the dominant signal, supporting its central role in coordinating early acclimation. Overall, these findings demonstrate that early salt acclimation in *L. maritima* is rapid but spatially and functionally uncoupled, combining efficient shoot osmotic stabilization with persistent photosynthetic constraints and increased root sensitivity.

## Supporting information

Dataset S1.Statistical analysis of all the comparisons performed in this study

Supplementary information. Includes Figures S1-4 and Tables S1 and S2.

## Acknowledgements

Special thanks to Rosa Padilla for her assistance in generating the leaf and root ionomic data. This work was funded by the Spanish Ministry of Science and Innovation (MICINN), grant number PID2019-104000-RB-I00 and Predoctoral Research Fellowship (PIF, Convocatòria 2021/D/LE/CC/3).

## Competing interests

None declared.

## Author contributions

RT and ML conceived the study and secured funding. CG, RT, ML and EB designed the methodology. CG, RT and ML prepared the first draft of the manuscript. CG and GCH collected the data. CG and ML analysed the data. ML, CG, RT and EB contributed critically to subsequent drafts. All authors approved the final version for publication.

## Supporting information

Dataset S1. Statistical analysis of all the comparisons performed in this study.

Fig. S1. Shoot ionomic profiles.

Fig. S2. Root ionomic profiles.

Fig. S3. Selected shoot element ratios.

Fig. S4. Root ionomic profiles.

Table S1. Primer list for transcript quantification.

Table S2. Root growth and architecture parameters.

## Notes

### Competing Interest Statement

The authors have declared no competing interest.

