## Supplementary information. Includes Figures S1-4 and Tables S1 and S2. for "Early salt shock uncouples shoot - root acclimation in *Lobularia maritima*"

### Supporting Information

Article title: Early salt shock uncouples shoot - root acclimation in *Lobularia maritima*
Authors: Carlos González-Cobo[1], Roser Tolrà[1], Gheith Chariff[1], Eliana Bianucci[1], Mercè Llugany[1]
Affiliations: [1]Plant Physiology (BABVE), Faculty of Biosciences, Universitat Autònoma de Barcelona, Cerdanyola del Vallès, 08193, Spain.

The following Supporting Information is available for this article:

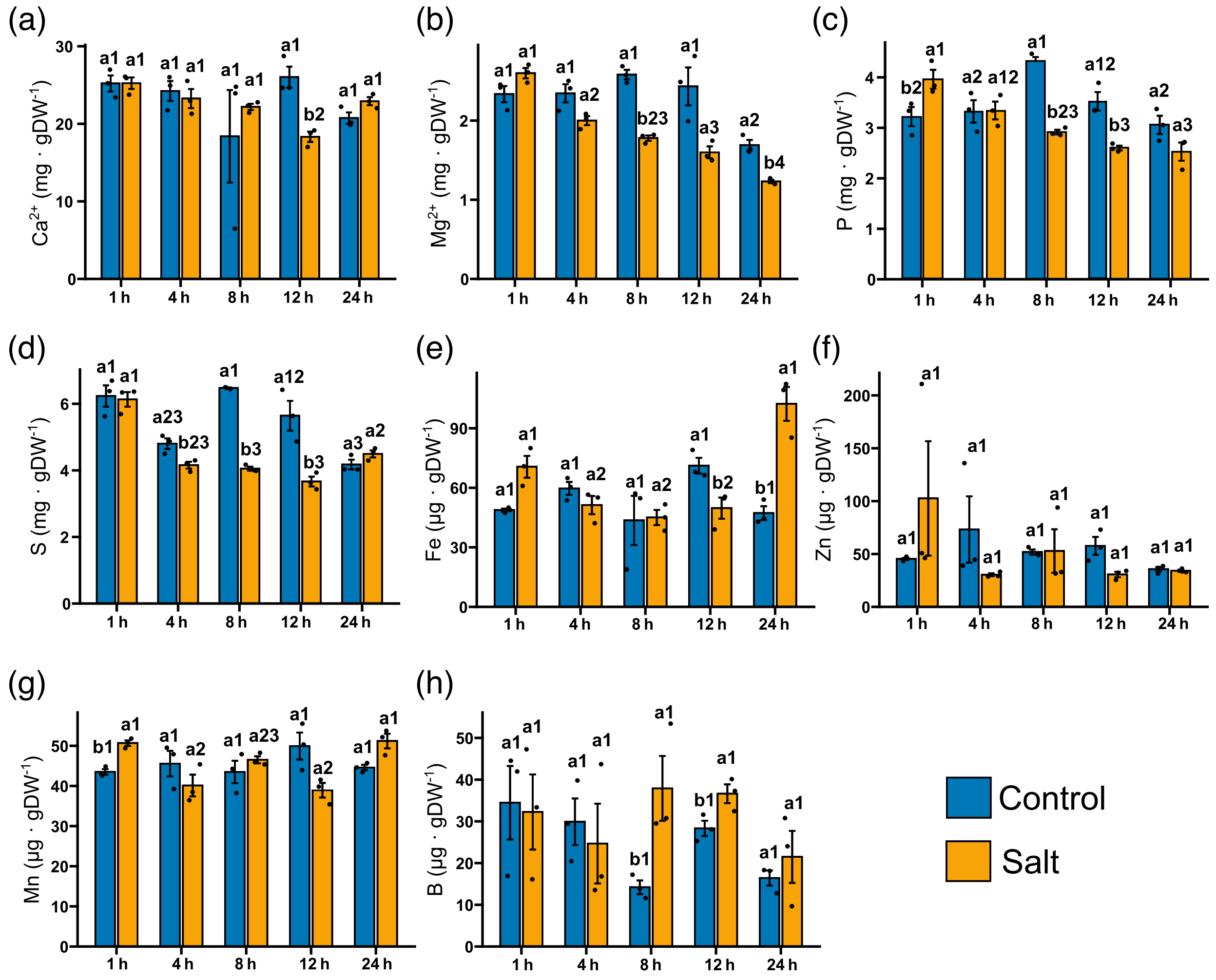


**Figure S1**. **Ionome profile of cultivar *L. maritima* leaves exposed to salt shock.** Barplots showing the mean ± standard error (SE) of (a) $Ca^{2+}$, (b) $Mg^{2+}$, (c) P, (d), S, (e) Fe, (f) Zn, (g) Mn, and (h) B shoot content. Plants were exposed to 0 mM NaCl (control) or 400 mM NaCl for the indicated time points. Different letters indicate significant differences between salt treatments (t-test, $p < 0.05$), while different numbers indicate significant differences among time points (t-test, $p < 0.05$).

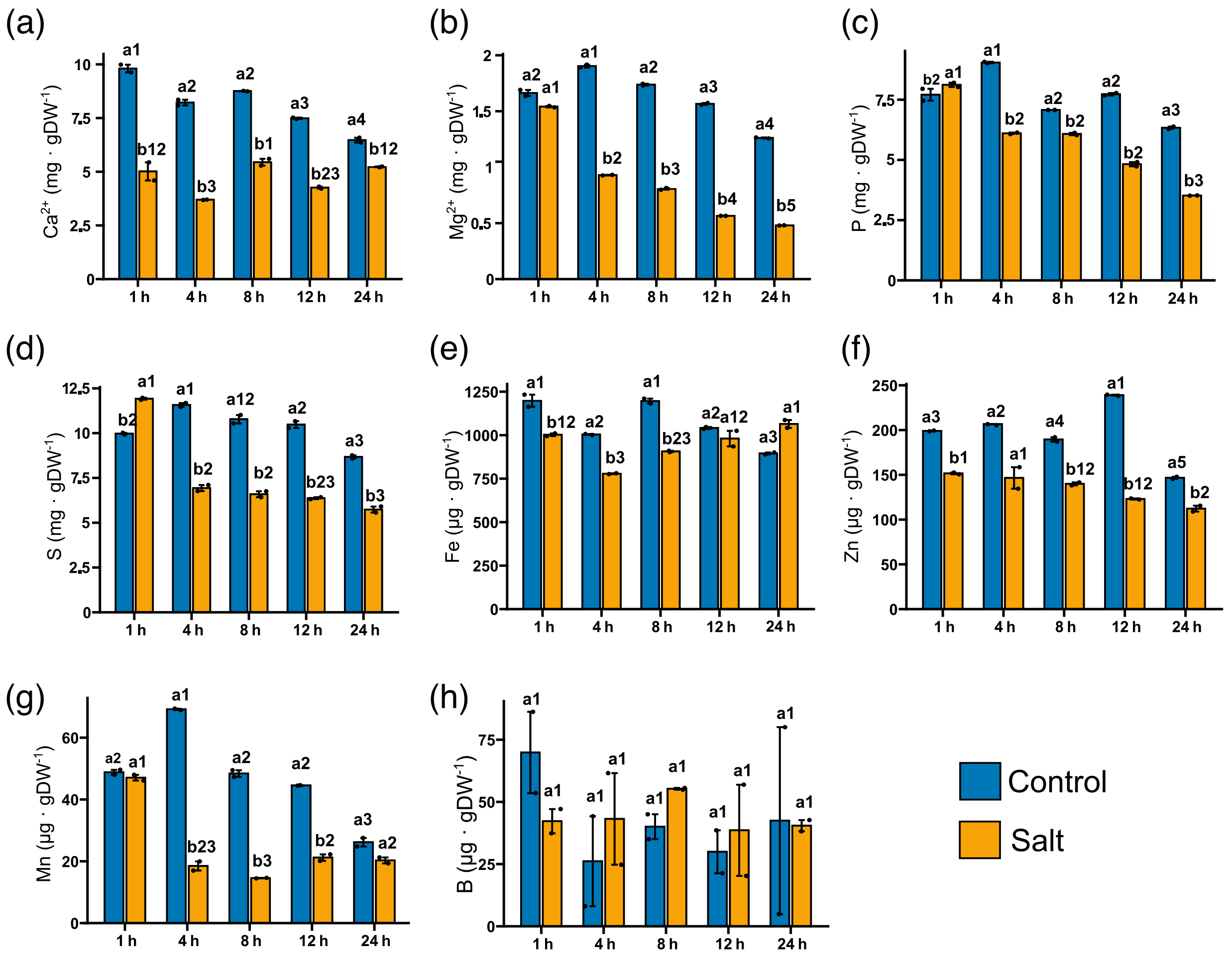


**Figure S2 Ionome profile of cultivar *L. maritima* roots exposed to salt shock.** Barplots showing the mean ± standard error (SE) of (a) $Ca^{2+}$, (b) $Mg^{2+}$, (c) P, (d), S, (e) Fe, (f) Zn, (g) Mn, and (h) B root content. Plants were exposed to 0 mM NaCl (control) or 400 mM NaCl for the indicated time points. Different letters indicate significant differences between salt treatments (t-test, $p < 0.05$), while different numbers indicate significant differences among time points (t-test, $p < 0.05$).

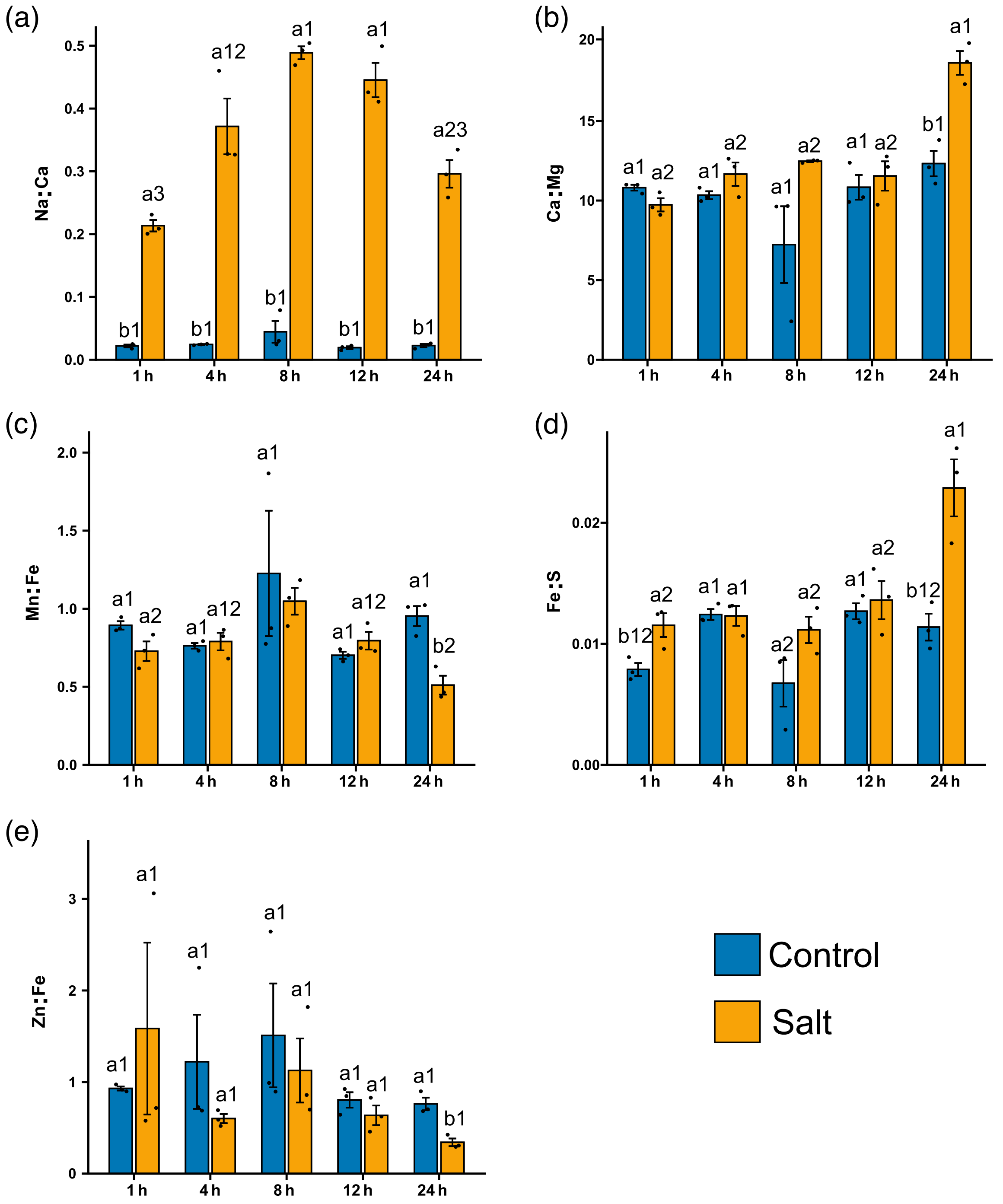


**Figure S3**. **Element ratios of cultivar *L. maritima* leaves exposed to salt shock**. Barplots showing the mean ± standard error (SE) of shoot (a) $Na^{+}$:$Ca^{2+}$, (b) $Ca^{2+}$:$Mg^{2+}$, (c) Mn:Fe, (d), Fe:S, and (e) Zn:Fe ratios. Plants were exposed to 0 mM NaCl (control) or 400 mM NaCl for the indicated time points. Different letters indicate significant differences between salt treatments (t-test, $p < 0.05$), while different numbers indicate significant differences among time points (t-test, $p < 0.05$).

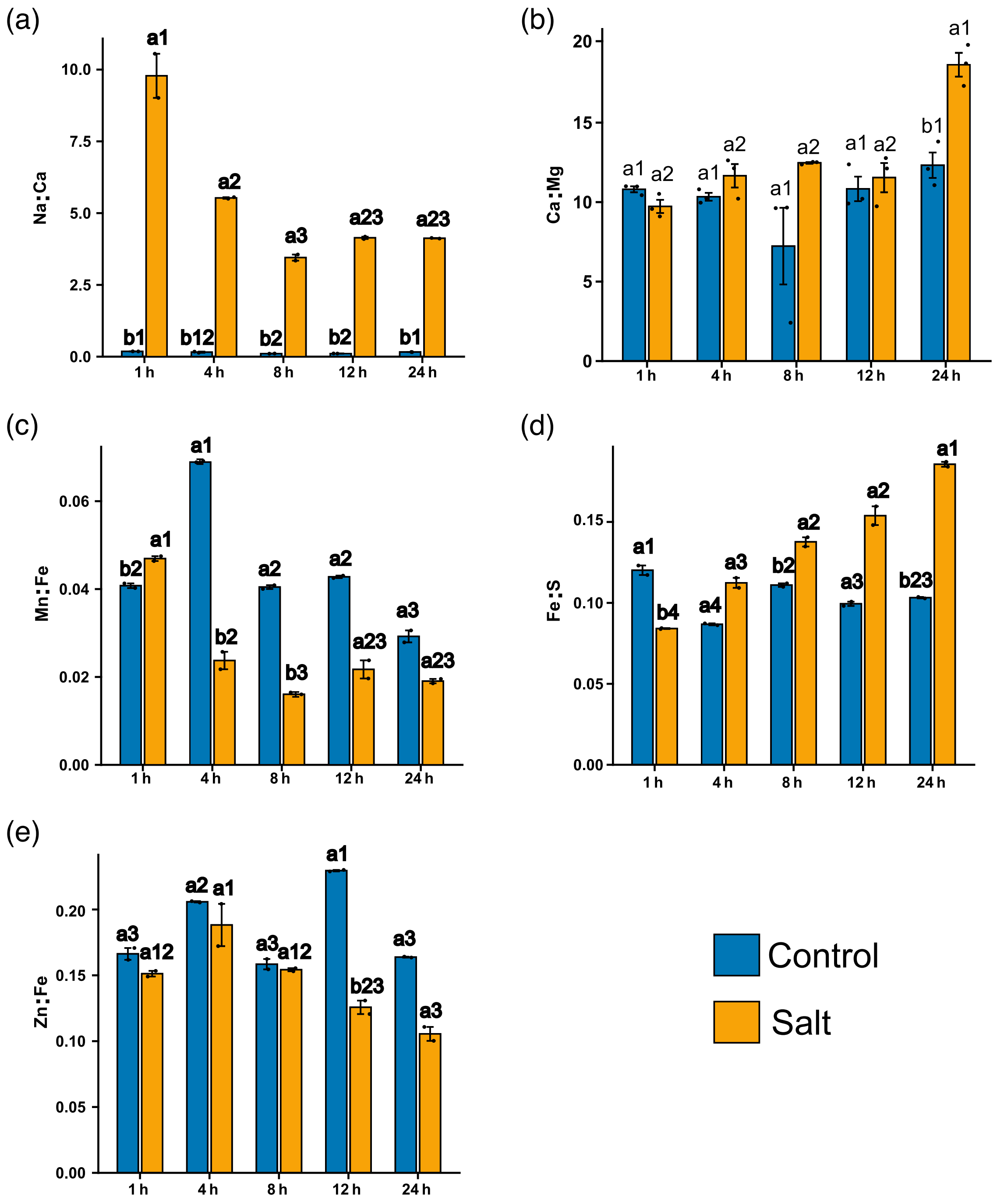


**Figure S4. Element ratios of cultivar *L. maritima* leaves exposed to salt shock**. Barplots showing the mean ± standard error (SE) of root (a) $Na^{+}$:$Ca^{2+}$, (b) $Ca^{2+}$:$Mg^{2+}$, (c) Mn:Fe, (d), Fe:S, and (e) Zn:Fe ratios. Plants were exposed to 0 mM NaCl (control) or 400 mM NaCl for the indicated time points. Different letters indicate significant differences between salt treatments (t-test, $p < 0.05$), while different numbers indicate significant differences among time points (t-test, $p < 0.05$).

**Table S1. Primers used for gene expression analysis of target genes.**

| Primer Name | Primer | Sequence (5’ -> 3’) |
|---|---|---|
| *HKT1_F* | Forward | CACTCTGTTCCCTTGCTTCT |
| *HKT1_R* | Reverse | ACGCTGTCATTCCAAGAATA |
| *NHK1_F* | Forward | CTGTGTGTACACTGCAGGTTC |
| *NHK1_R* | Reverse | CCTCGTGGTTAAGGTGTGTG |
| *SOS1_F* | Forward | AGTCCGATCTTCACTTCCTC |
| *SOS1_R* | Reverse | CTTCAAGCCAGTCTGGACAG |
| *SOS2_F* | Forward | CGGACTCAGTGCATTGCCTC |
| *SOS2_R* | Reverse | AACATCTGCTGCTGAACCGT |
| *SOS3_F* | Forward | CGAGCGCACCTGTCCATGAC |
| *SOS3_R* | Reverse | TTGCGGTCAGCTTCGATAAA |
| *UBQ10_F* | Forward | ACACCATCGACAACGTCAAG |
| *UBQ10_R* | Reverse | TACCACCACGGAGCCTGAGC |

**Table S2. Root developmental analysis of cultivar *L. maritima* seedlings exposed to salt shock.** Mean ± standard error (SE) of selected root morphological parameters obtained using WinRhizo software. Plants were exposed to 0 mM NaCl (control) or 400 mM NaCl for the indicated time points. Different letters indicate significant differences between salt treatments (t-test, p <0.05), while different numbers indicate significant differences among time points (Tukey HSD, p <0.05).

| Time point | Treatment | Root length (cm) | Root volume ($cm^3$) | Forks | Tips |
|---|---|---|---|---|---|
| 4h | Control | 73.77 ± 6.10 a1 | 0.1506 ± 0.028 a2 | 284.6 ± 37.0 a1 | 62.0 ± 6.67 a2 |
| | Salt | 53.98 ± 6.63 a1 | 0.1448 ± 0.011 a1 | 248.2 ± 50.8 a1 | 55.6 ± 10.59 a1 |
| 8h | Control | 64.18 ± 2.72 a1 | 0.1435 ± 0.008 a2 | 262.8 ± 15.1 a1 | 125.0 ± 9.30 a1 |
| | Salt | 58.81 ± 5.13 a1 | 0.1602 ± 0.008 a1 | 269.6 ± 26.3 a1 | 52.6 ± 5.82 b1 |
| 12h | Control | 55.67 ± 7.81 a1 | 0.1494 ± 0.013 a2 | 278.6 ± 34.3 a1 | 77.8 ± 9.98 a2 |
| | Salt | 50.58 ± 10.37 a1 | 0.1980 ± 0.016 a1 | 244.2 ± 45.6 a1 | 68.2 ± 16.30 a1 |
| 24h | Control | 60.68 ± 5.46 a1 | 0.2338 ± 0.018 a1 | 307.5 ± 35.0 a1 | 67.0 ± 9.90 a2 |
| | Salt | 39.48 ± 3.47 b1 | 0.1860 ± 0.025 a1 | 185.0 ± 14.7 b1 | 49.3 ± 3.54 a1 |
